# Brainstem Neurons that Command Left/Right Locomotor Asymmetries

**DOI:** 10.1101/754812

**Authors:** Jared M. Cregg, Roberto Leiras, Alexia Montalant, Ian R. Wickersham, Ole Kiehn

## Abstract

Descending command neurons instruct spinal networks to execute basic locomotor functions, such as which gait and what speed. The command functions for gait and speed are symmetric, implying that a separate unknown system directs asymmetric movements—the ability to move left or right. Here we report the discovery that *Chx10*-lineage reticulospinal neurons act to control the direction of locomotor movements in mammals. *Chx10* neurons exhibit ipsilateral projection, and can decrease spinal limb-based locomotor activity ipsilaterally. This circuit mechanism acts as the basis for left or right locomotor movements in freely moving animals: selective unilateral activation of *Chx10* neurons causes ipsilateral movements whereas inhibition causes contralateral movements. Spontaneous forward locomotion is thus transformed into an ipsilateral movement by braking locomotion on the ipsilateral side. We identify sensorimotor brain regions that project onto *Chx10* reticulospinal neurons, and demonstrate that their unilateral activation can impart left/right directional commands. Together these data identify the descending motor system which commands left/right locomotor asymmetries in mammals.

## INTRODUCTION

Locomotion is a natural behavior universal to the animal kingdom. In vertebrates, coordination of rhythmic locomotor movements occurs largely within circuits of the spinal cord itself (1–7). For these circuits to function, they need commands from supraspinal effector neurons that control the start and speed of locomotion. The brainstem command neurons which control these parameters have been examined extensively in several vertebrate species (8–20). Recently, brainstem neurons that mediate locomotor stop were also identified (9, 11, 15).

Characteristically, when start command pathways of the midbrain locomotor region are activated unilaterally, they cause bilateral full-bodied locomotion which proceeds in a straight line (10, 11, 14, 17, 18, 20–23). This finding underscores symmetry in the command for initiating locomotion and controlling its speed. The anatomical basis for this symmetry has been worked out in some detail. At the level of the midbrain locomotor region, which is comprised of glutamatergic neurons in the cuneiform and pedunculopontine nuclei, neurons exhibit extensive commissural connectivity with neurons of the contralateral side (10, 19). The start command is then relayed to reticulospinal neurons, including those of the lateral paragigantocellular nucleus (LPGi), which in turn activate spinal locomotor circuits (11, 21, 24– 29). These reticulospinal neurons receive bilateral projections from the midbrain locomotor region (10, 11, 21, 23). Neurons of the LPGi project bilaterally, innervating both sides of the spinal cord, and unilateral optogenetic activation of LPGi glutamatergic neurons initiates symmetric full-bodied locomotion (11). Unilateral stimulation of neurons in the parapyramidal area also causes symmetric locomotor activity in (16).

A considerable gap in our knowledge is the inability to explain how command neurons direct locomotor movements to the left or right side (30). A system which executes left/right locomotor asymmetries would be required for any goal-directed locomotor movement, as might occur during basic behaviors like foraging, navigation, and/or escape, but also during specialized locomotor tasks (31). From *in vitro* studies, it is clear that rhythmogenic modules can operate independently within the left or right spinal cord in mammals (32–38). Nonetheless, differential engagement of left/right rhythmogenic modules cannot be mediated by a symmetric start command. Moreover, unilateral lesion of the corticospinal tract (i.e. unilateral pyramidotomy) does not result in any overt left/right locomotor asymmetries (39, 40). Although unilateral labyrinth ablation or vestibulocochlear nerve (VIII) stimulation can initiate reflexive rotational behavior (41), this self-righting reflex is not a voluntary locomotor command.

In the present work, we hypothesized that a turn in limbed animals could be implemented by inhibition of locomotor circuits on one side. We focused on a system of nucleus gigantocellularis (Gi) reticulospinal neurons uniquely identified by expression of *Chx10* (9). These *Chx10* ‘stop neurons’ are glutamatergic, and their bilateral activation arrests locomotion by suppressing locomotor rhythmogenesis in the spinal cord (9). We found that *Chx10* Gi neurons 1) exhibit dominant unilateral projection to the spinal cord that can bias locomotor activity unilaterally, 2) define the direction of locomotion by effecting changes in ipsilateral limb and axial movements *in vivo*, and 3) can be engaged to impart asymmetric movements via unilateral input from distinct sensorimotor brain regions. *Chx10* Gi neurons exhibit all the features of a *bona fide* system for executing left/right locomotor behaviors in limbed animals.

## RESULTS

### *Chx10* Gi neurons form a prominent spinal tract of ipsilaterally projecting axons

For *Chx10* Gi neurons to regulate locomotion unilaterally, they should exhibit predominant unilateral projection to the spinal cord. We examined this by labeling *Chx10*-lineage neurons of the rostral gigantocellularis with an anterograde viral tracer (*Chx10*^*Cre*^ > AAV-FLEX-tdTomato-2A-synGFP) (Figure 1A). In adult mice, a unilateral injection labeled *Chx10* Gi neurons predominantly on the same side (Figures 1B-1D). *Chx10* neurons projected axons caudally (Figures S1A and S1B), which then coalesced to form a prominent ipsilateral tract of axons lateral to the inferior olive (Figure S1C). At the level of the pyramidal decussation, this tract of *Chx10* reticulospinal axons turned ventrally to occupy the ipsilateral ventral funiculus within the rostral-most segment of the spinal cord (Figure S1C). TdTomato^+^ axons arborized predominately within the ipsilateral cord (Figures 1E and 1F), although some arborizations could also be found on the contralateral side.

**Figure 1.**
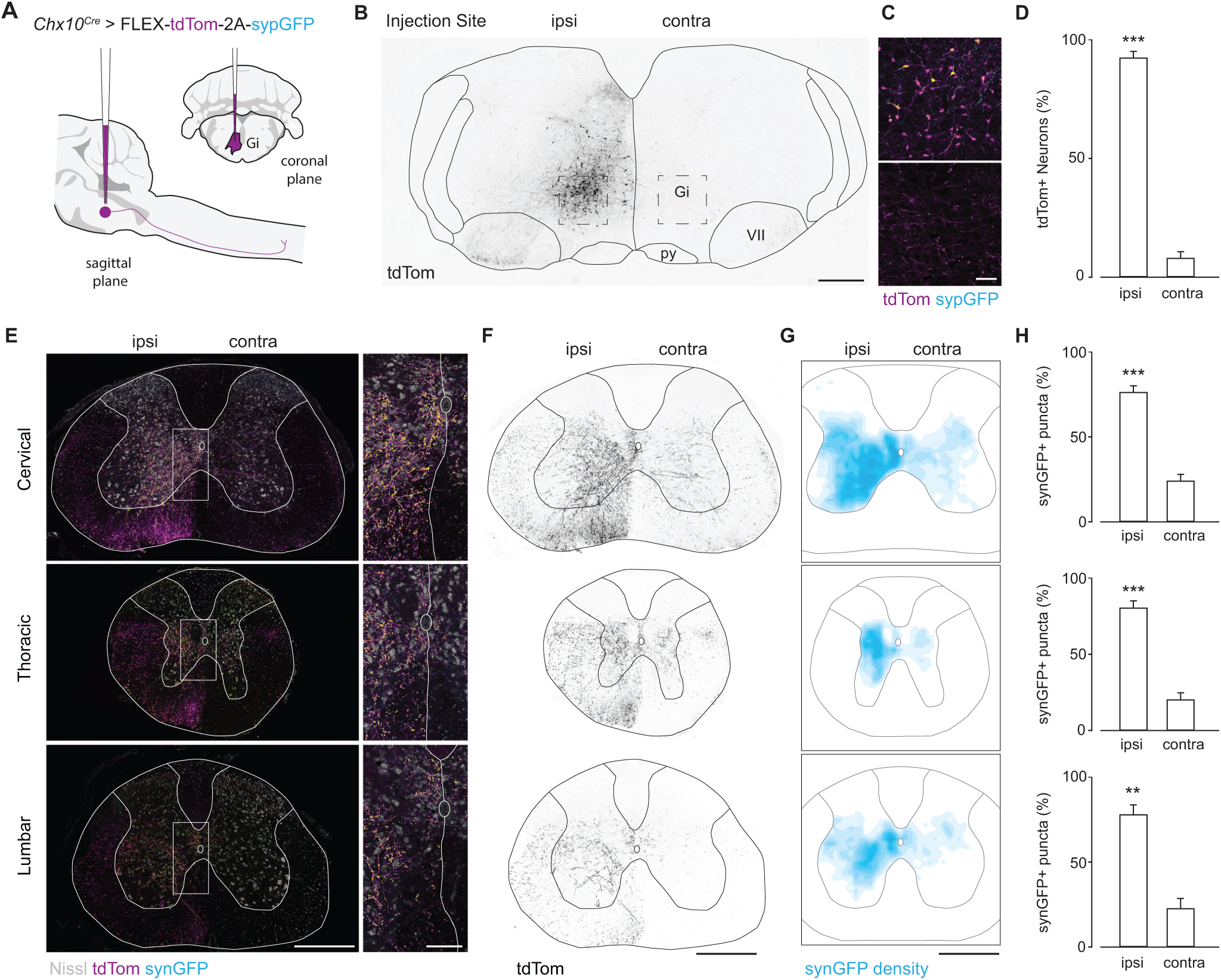
*Chx10* Gi neurons form a prominent tract of descending axons that project ipsilaterally. **A**, Unilateral labeling of *Chx10* neurons of the rostral gigantocellularis (Gi) using the Cre-dependent anterograde tracer AAV-FLEX-tdTom-2A-synGFP. **B**, Inverted fluorescent image of tdTomato^+^ neurons at the injection site within Gi. **C**, *Top*, tdTom^+^/synGFP^+^ neurons at injection site. *Bottom*, sparse tdTom^+^ axonal labeling and synGFP^+^ punctae contralateral to the injection site. Scale bar = 50 µm. **D**, Quantification of tdTom^+^ neurons labeled ipsilateral (ipsi) and contralateral (contra) to the injection site. \*\*\**P* = 3×10^−5^, two-tailed t-test, *n* = 3. **E**, *Left*, A prominent tract of tdTom^+^ axons is observed within the ventrolateral funiculus of the cervical (top), thoracic (middle), and lumbar (bottom) spinal cord. Axons project almost exclusively on the ipsilateral side. SynGFP^+^ punctae (overlap of cyan with magenta yields yellow) exhibit a high density on the ipsilateral side. Scale bar = 500 µm. *Right*, Insets from images on left demonstrate a sharp division in the density of axon profiles and synGFP^+^ punctae between the ipsilateral and contralateral sides. Scale bar = 100 µm. **F**, Inverted fluorescent images of tdTom^+^ axonal projections in the cervical (top), thoracic (middle), and lumbar (bottom) spinal cord. Axons descend in the ventrolateral funiculus within a well-defined “wedge”. Scale bar = 500 µm. **G**, Density plots for synGFP^+^ punctae within the gray matter of the cervical (top), thoracic (middle), and lumbar (bottom) spinal cord (average from 3 animals). The density of putative synapses is greatest in the ipsilateral intermediate gray matter—laminae VII, VIII, and X. SynGFP^+^ punctae were excluded from laminae I-VI of the dorsal horn, lamina IX (motor neurons), and at the thoracic level, Clarke’s column. Scale bar = 500 µm. **H**, Quantification of synGFP^+^ punctae within the ipsilateral and contralateral spinal cord gray matter. Cervical, \*\*\**P* = 0.0006; Thoracic, \*\*\**P* = 0.0002; Lumbar, \*\**P* = 0.007. Two-tailed t-test, *n* = 3.

We quantified the position and density of *Chx10* Gi syn-GFP+ punctae (putative synapses) within the spinal cord. A vast majority of synGFP^+^ punctae were ipsilateral to the injection site in the cervical, thoracic, and lumbar cord (76.3-80.6%, Figures 1G and 1H), with fewer synGFP^+^ punctae on the contralateral side. The density of synGFP^+^ punctae was greatest in the ipsilateral intermediate gray (laminae VII, VIII, and X) (Figures 1G and 1H), with the exception of Clarke’s column (Th1 to L3) (Figures 1G and 1H), a medial nucleus which conveys proprioceptive inputs to the cerebellum. Notably, synGFP^+^ punctae were largely absent from dorsal horn laminae I-VI where sensory networks are localized, and from lamina IX where motor neurons reside. These data indicate that *Chx10* reticulospinal neurons likely act on rhythmic premotor networks (7, 9), rather than on motor neurons themselves (9). We conclude that *Chx10* Gi neurons have predominantly ipsilateral projections to the spinal cord, a feature that may allow for differential control of left and right spinal motor networks.

### *Chx10* Gi neurons enable differential control of rhythmic motor activity on the left and right sides

We next asked whether ipsilateral projection of *Chx10* Gi neurons can allow for differential control of left and right spinal motor networks using an *in vitro* split-bath preparation (Figure 2A). Here, locomotor-like activity is maintained in the spinal cord and synaptic activity is blocked in the brainstem. Stimulating *Chx10* Gi neurons in this configuration excludes the possibility that locomotor effects are due to axon collaterals in the brainstem (9, 42). In *Chx10*^*Cre*^;*R26R*^*ChR2*^ preparations, we found that unilateral stimulation of *Chx10* Gi neurons with blue light could arrest rhythmic hindlimb locomotor activity only on the side ipsilateral to stimulation with little effect on the rhythm of the contralateral side (Figure 2B, *n* = 2/4). In other examples, we found a decrease in the amplitude of locomotor bursting ipsilateral to photo-stimulation (Figure 2C). Altogether, burst amplitude exhibited a 57 ± 12% reduction ipsilateral (but not contralateral) to photostimulation (Figure 2D). These observations phenocopy data demonstrating that rhythmicity can be initiated on one side of the cord (34, 43), and that unilateral stimulation of inhibitory neurons in the lumbar spinal cord can slow or arrest locomotor rhythmogenesis only on the side of stimulation (34). We conclude that *Chx10* reticulospinal neurons can function as a unilateral locomotor effector.

**Figure 2.**
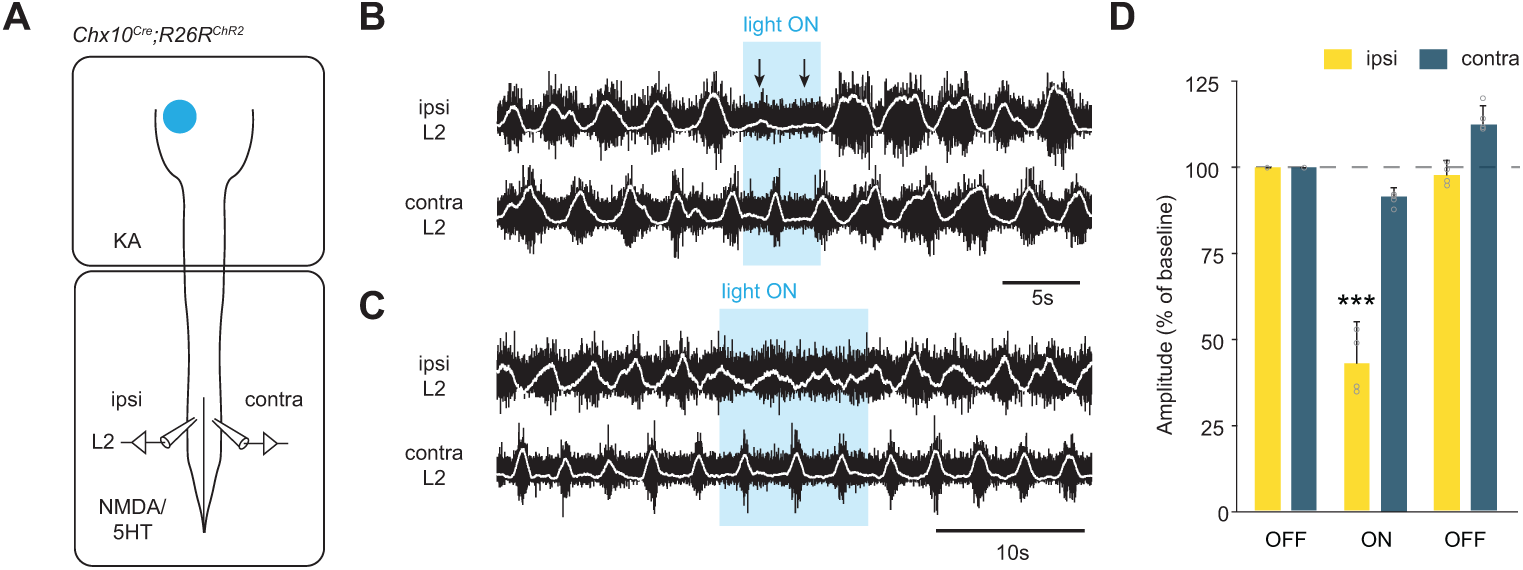
*Chx10* Gi neurons can arrest and/or decrease *in vitro* locomotor-like activity on the same side. **A**, Schematic of split-bath brainstem-spinal cord preparation from P0-2 *Chx10*^*Cre*^;*R26R*^*ChR2*^ mice, which was used to interrogate unilateral function of *Chx10* reticulospinal neurons *in vitro*. A rostral brainstem compartment was bathed with kynurenic acid (4 mM) to block all glutamatergic transmission amongst axon collaterals (42), and a caudal spinal cord compartment was bathed with 5HT (8 µM) and NMDA (8 µM) to induce locomotor-like activity, assayed by recording from the L2 ventral roots. **B**, Unilateral photostimulation of *Chx10* Gi neurons can arrest locomotor-like activity ipsilateral (but not contralateral) to stimulation. **C**, In another example, unilateral photostimulation of *Chx10* Gi neurons decreases the amplitude of motor bursts ipsilateral (but not contralateral) to stimulation. **D**, Quantification of burst amplitude before, during, and after photostimulation. \*\*\**P* <1.7×10^−4^, one-way ANOVA, *n* = 4.

### Unilateral excitation of *Chx10* Gi neurons *in vivo* reveals a role in control of left/right movements

The capacity of *Chx10* Gi neurons to act as an ipsilateral locomotor effector *in vitro* suggests a function of *Chx10* Gi neurons in controlling locomotor asymmetries—the ability to move left or right. To test this, we examined the behavioral consequence of unilateral stimulation of *Chx10* Gi neurons in freely moving mice. We expressed excitatory hM3Dq-DREADDs in *Chx10* Gi neurons on one side of the brainstem (Figure 3A). hM3Dq-DREADDs can be activated with low doses (0.5 mg kg^−1^) of clozapine-N-oxide (CNO), which causes neuronal depolarization via G_q_-mediated signaling (44). In a cylinder assay, which promotes locomotor turning, we found that CNO administration caused a strong preference in turning toward the side of *Chx10* Gi activation (the ipsilateral side, Figure 3B). We next examined locomotion in an open field. CNO administration strongly induced ipsilateral turning even during unrestricted, spontaneous locomotion (Figure 3C, Video S1). CNO administration had no behavioral effect in control injected animals (Figure S2).

**Figure 3.**
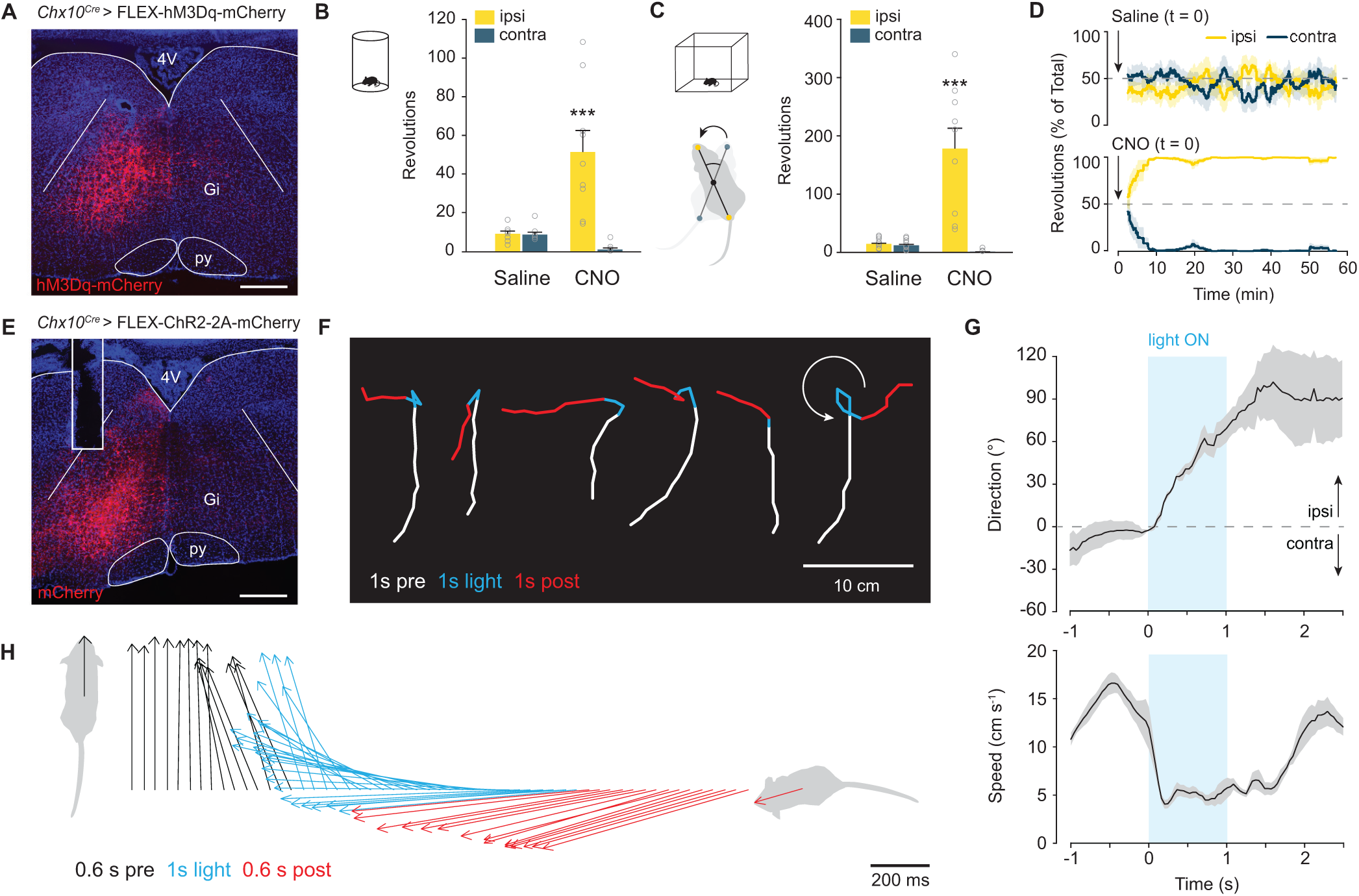
Excitation of *Chx10* Gi neurons causes ipsilateral movements. **A**, Unilateral infection of *Chx10* Gi neurons with AAV-FLEX-hM3Dq-mCherry. Scale bar = 500 µm. **B**, Movement preference in a 10 minute cylinder assay 1 h after injection of saline or CNO. \*\*\**P* < 4×10^−5^, one-way ANOVA with Tukey HSD, *n* = 9. **C**, Movement preference in an open field assay, quantified between 30-60 min post-injection of saline or CNO. \*\*\**P* < 3×10^−6^, one-way ANOVA with Tukey HSD, *n* = 9. **D**, Instantaneous analysis of movement preference, quantified as the percentage of ipsilateral versus contralateral revolutions (bin = 5 min), following injection of saline (*top*) or CNO (*bottom*) at t = 0. **E**, Unilateral injection of AAV-FLEX-ChR2-2A-mCherry in *Chx10*^*Cre*^ mice. Example of mCherry expression and optical fiber placement in Gi. Scale bar = 500 µm. **F**, Body tracking 1 s before (white), 1 s during (blue), and 1 s following (red) light stimulation. Examples (left) are from 5 different animals. The example on the right demonstrates a 270 degree turn. The distance traveled during stimulation is reduced compared to before and after stimulation. **G**, Quantification of movement direction and speed relative to light onset. Photostimulation causes an abrupt shift in movement direction toward the ipsilateral side, accompanied by a reduction in locomotor speed. Data are mean ± standard error mean, *n* = 5, 3-9 trials per animal. **H**, Body center-to-head vectors are plotted as a function of time 0.6 s before (black), 1 s during (blue), and 0.6 s following (red) light stimulation.

Quantification of the percentage of ipsilateral versus contralateral revolutions in open-field analysis revealed that ipsilateral turning developed over time (Figure 3D), approaching 100% ipsilateral turning preference by 10 minutes. At early stages following drug administration (5-15 minutes), these ipsilateral turns were smooth—similar to those observed during spontaneous changes in locomotor direction. There were no clear changes in trunk (axial) posture at early stages following administration of CNO. At late stages (> 20 min), we observed an ipsilateral axial bend which was evident even at rest (Video S1). Notably, during both early and late stages following administration of CNO, *bona fide* turns—where locomotor direction actually changed—only occurred when the animal started moving forward, suggesting specific involvement of the limbs in imparting changes to locomotor direction.

To further investigate the dynamics of this behavior, we performed short-lasting photo-stimulation of *Chx10* Gi neurons after unilateral expression of channelrhodopsin-2 (*Chx10*^*Cre*^ > AAV-FLEX-ChR2) (Figure 3E). Animals were stimulated with blue light (473 nm) for 1 s when slowly walking in an open field. Movement trajectories were calculated 1 s before, during, and after stimulation (Figure 3F). In all cases (27 trials in 5 animals), photostimulation led to an abrupt turn toward the ipsilateral side (Figure 3F, Video S2). This ipsilateral turn was initiated within 150 ms and lasted approximately 1.2 s (Figure 3G and 3H), and was temporally correlated with a reduction in locomotor speed (Figure 3F and 3G). Photostimulation at rest could not evoke a locomotor turn, and instead caused head and trunk bending toward the ipsilateral side (Video S2). In some cases, the animal would perform a 270-degree turn (Figure 3F). Such sharp turns were accompanied by bending of the trunk, suggesting involvement of the axial muscles at least for pronounced changes in direction.

These experiments indicate that *Chx10* Gi neurons evoke ipsilateral movements via two distinct mechanisms. Stimulation of *Chx10* Gi neurons at rest evoked ipsilateral trunk/head movements, whereas stimulation during ongoing locomotion caused locomotor movements toward the ipsilateral side. *Chx10* Gi neurons appear to encompass all modes of turning; increasing *Chx10* Gi activity caused changes in trunk posture at rest as well as either gradual and/or sharp changes in locomotor direction.

### Inhibition of *Chx10* Gi neurons causes contralateral movements

We next examined the behavioral consequence of unilateral inactivation of *Chx10* Gi neurons using a Cre-dependent tetanus toxin virus (*Chx10*^*Cre*^ > AAV-FLEX-TeLC-GFP) (Figure 4A). In the cylinder assay, we found that FLEX-TeLC (but not FLEX-GFP) injected mice exhibited a strong turning preference toward the contralateral side (Figure 4B). We performed a time course analysis of turning preference after TeLC injection in the open field. One day before injection of FLEX-TeLC-GFP or FLEX-GFP, mice exhibited no open-field turning preference (Figures 4C and 4D, Video S3). Animals injected unilaterally with FLEX-TeLC (but not FLEX-GFP) started to exhibit turning contralateral to the injected side as soon as 3 days after injection (Figures 4C and 4D). By 9 days, 100% of TeLC animals exhibited contralateral turning (Figures 4C and 4D, Video S3). At days 3-6, contralateral turns were smooth, and there were no obvious effects on trunk posture when the animal was at rest (Video S3). When fully developed at day 9, turns were sharper than in the early stages (Video S3). Therefore, similar to unilateral *Chx10* Gi stimulation, inhibition of *Chx10* Gi activity caused both gradual and sharp modes of locomotor turning–but toward the contralateral side. We further confirmed these results by acute inactivation of *Chx10* Gi neurons using inhibitory DREADDS (*Chx10*^*Cre*^ > FLEX-hM4Di), which also promoted contralateral movements (Figures 4E-4H).

**Figure 4.**
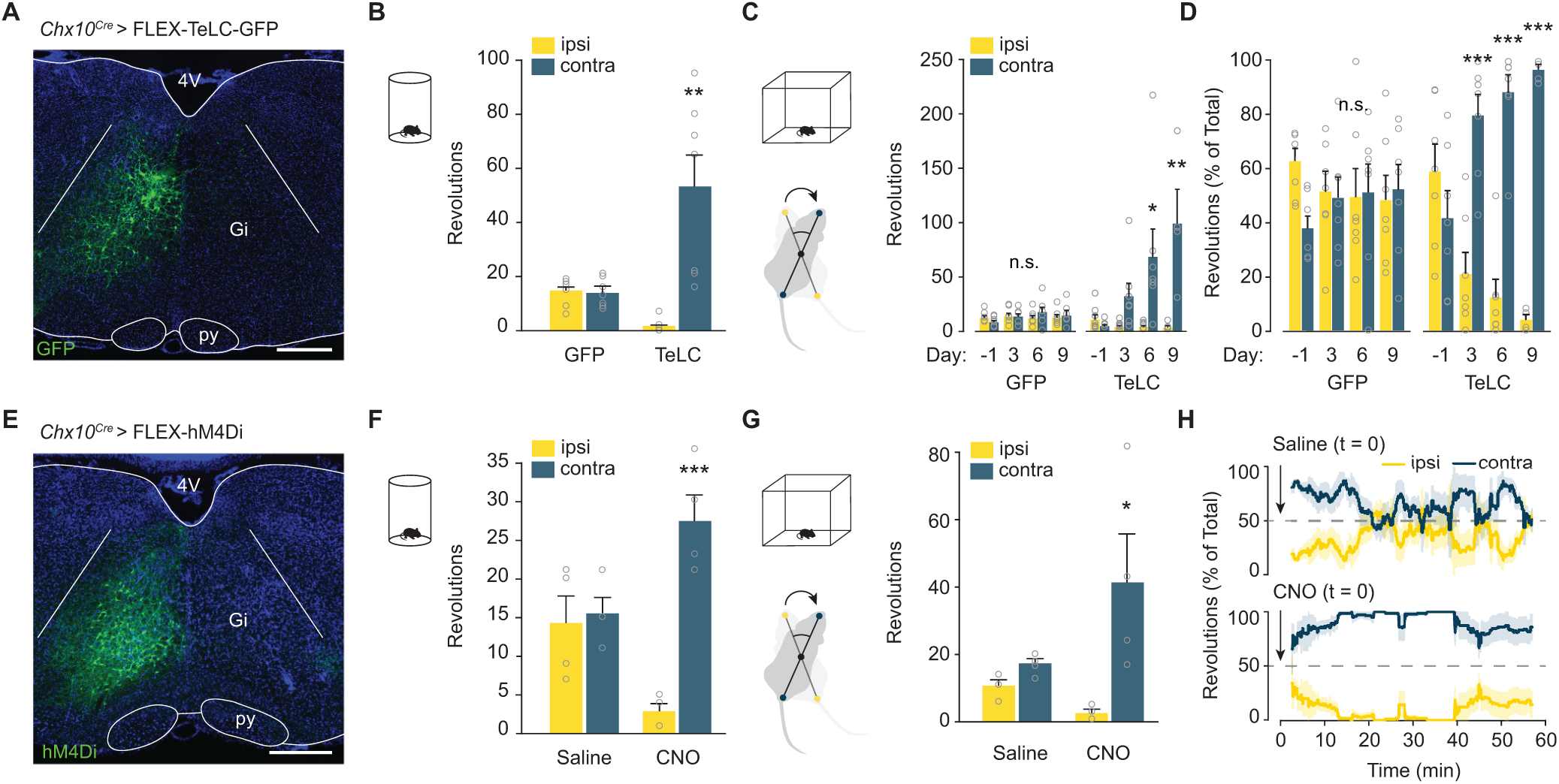
Inhibition of *Chx10* Gi neurons causes contralateral movements. **A**, Unilateral injection of AAV-FLEX-TeLC-GFP or AAV-FLEX-GFP in *Chx10*^*Cre*^ mice. Example of GFP expression in Gi 2 weeks post-injection. Scale bar = 500 µm. **B**, Turning preference in a 10 minute cylinder assay 7 d after viral injection. \*\**P* < 0.0013, one-way ANOVA with Tukey HSD, *n* = 7. **C**, Turning preference in an open field assay, quantified as the total number of revolutions. GFP: *P* > 0.35 for each comparison. TeLC: day -1, *P* = 0.99 ipsi vs. contra; day 3, *P* = 0.77; day 6, \**P* = 0.017; day 9, \*\**P* = 0.0048. One-way ANOVA with Tukey HSD. **D**, Turning preference in an open field assay quantified as the percentage of total revolutions over a 30 min trial. GFP: *P* > 0.41 for each comparison. TeLC: day -1, *P* = 0.76 ipsi vs. contra; day 3, \*\*\**P* = 9.7×10^−5^; day 6, \*\*\**P* = 5.9×10^−7^; day 9, \*\*\**P* = 3.2×10^−6^. One-way ANOVA with Tukey HSD. **E**, Example of hM4Di expression after unilateral injection of AAV-FLEX-hM4Di in *Chx10*^*Cre*^ mice. **F**, Movement preference in a 10 minute cylinder assay 20 min after injection of saline or CNO. \*\*\**P* = 0.0002, one-way ANOVA with Tukey HSD, *n* = 4. **G**, Movement preference in an open field assay, quantified between 30-60 min post-injection of saline or CNO. \**P* = 0.012, one-way ANOVA with Tukey HSD, *n* = 4. **H**, Instantaneous analysis of movement preference, quantified as the percentage of ipsilateral versus contralateral revolutions (bin = 5 min), following injection of saline (top) or CNO (bottom) at t = 0.

Unilateral inactivation of *Chx10* Gi neurons dramatically increased the total distance animals moved during a 30 min probe (Figure S3). These data suggest that *Chx10* Gi neurons act as a unilateral ‘brake’ on locomotion, and removing this unilateral brake dramatically increases locomotor speed.

### Limb dynamics during natural and drug-evoked turns

Our data strongly suggest that the mechanism for *Chx10*-evoked locomotor turns reflects distinct changes in limb dynamics: *Chx10*-evoked ipsilateral turns were accompanied by a decrease in locomotor speed (Figure 3F and 3G), whereas inhibition of *Chx10* Gi neurons caused contralateral turning accompanied by an increase in overall locomotor speed (Figure S3).

To investigate these limb dynamics in greater detail, we analyzed how limb coordination relates to locomotor direction in freely moving mice. DeepLabCut was used for markerless extraction of forelimb and hindlimb paw position in an open field (Video S4) (45). We focused on continuous locomotor bouts with a speed >15 cm s^−1^, which corresponds to fast walk or trot (46). For locomotor bouts in wild-type animals, locomotion that proceeded in a straight line exhibited a similar stride length on the left and right sides (Figure 5A). During spontaneous turns, stride length was reduced on the side of the turn (Figure 5A). We compared spontaneous turns in wild-type mice with those biased to one side via excitation (*Chx10*^*Cre*^ > hM3Dq-DREADDs after CNO) or inhibition (*Chx10*^*Cre*^ > FLEX-TeLC) of *Chx10* Gi neurons (Figures 5B and 5C). Turns in wild-type animals and those caused by *Chx10*^*Cre*^ > FLEX-hM3Dq or *Chx10*^*Cre*^ > FLEX-TeLC shared a defining characteristic, that is, the hindlimbs and the forelimbs on the side of the turn travelled a shorter distance than the leg opposite to the turn (Figures 5B and 5C).

**Figure 5.**
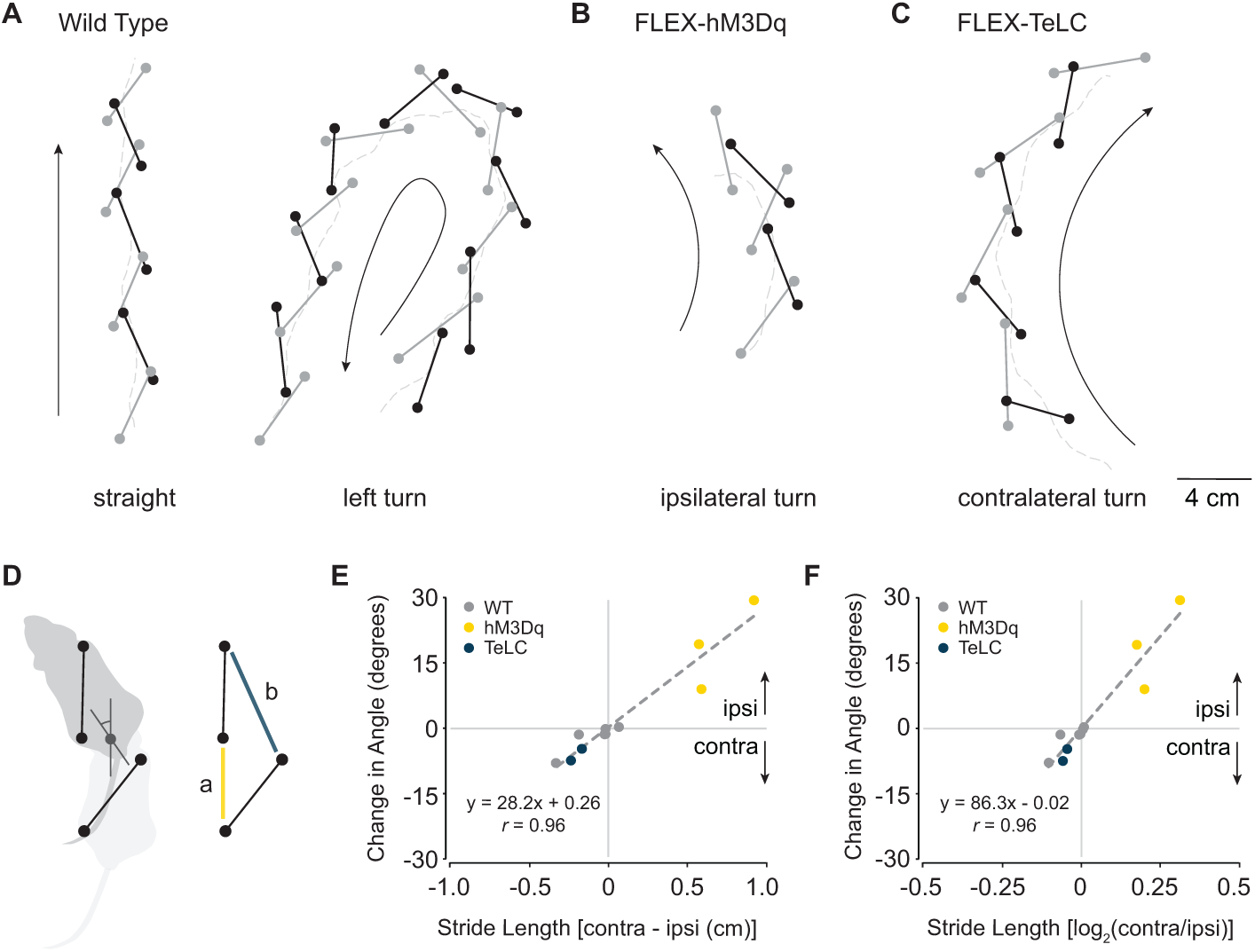
Limb dynamics during natural and *Chx10*-induced turns. **A**, Examples of footfalls in a wild-type mouse walking straight and making a spontaneous left turn. Black lines indicate the left-forelimb/right-hindlimb diagonal, whereas gray lines indicate the right-forelimb/left-hindlimb diagonal. For spontaneous turns, the limb on the side of the turn exhibits shorter steps than its corresponding diagonal. **B**, Example of footfalls in a left Gi *Chx10*^*Cre*^ > FLEX-hM3Dq mouse after injection of CNO, which causes turning toward the ipsilateral (left) side. Steps are shorter on the ipsilateral side. **C**, Example of footfalls in a left Gi *Chx10*^*Cre*^ > FLEX-TeLC mouse, which causes turning toward the contralateral (right) side. Steps are longer on the ipsilateral side. **D**, Schematic of turning based on data in A-C. A single step can exhibit left/right motor asymmetry, where the ipsilateral hindlimb exhibits a shorter stride length (a) than the diagonal limb of the contralateral side (b) leading to changes in heading position. **E-F** Mathematical models demonstrating that a difference in stride length predicts a difference in heading position. We quantified footfalls in wild-type, *Chx10*^*Cre*^ > FLEX-hM3Dq, and *Chx10*^*Cre*^ > FLEX-TeLC mice. For wild-type mice, the difference in stride length, measured either as the absolute distance (E) or as the log_2_ ratio (F), hovered around zero, which corresponds to straight walking and/or slight turning to one side. For FLEX-hM3Dq mice after CNO injection, the values are positive—predicting an ipsilateral turn. For FLEX-TeLC mice, the values are negative—predicting a contralateral turn.

We next asked whether differences in stride length between the left and right sides were sufficient to predict locomotor direction (Figure 5D). We calculated the relationship between the distance travelled by the right and left limbs with heading direction for all locomotor bouts sampled in wildtype, *Chx10*^*Cre*^ > FLEX-hM3Dq, and *Chx10*^*Cre*^ > FLEX-TeLC mice. In wild-type mice, we observed epochs with no difference in left/right stride length, which corresponded to straight walking, as well as epochs with longer left leg steps (turning right) and longer right leg steps (turning left). *Chx10*^*Cre*^ > FLEX-hM3Dq mice exhibited shorter steps on the side of injection, which caused ipsilateral turns. *Chx10*^*Cre*^ > FLEX-TeLC mice exhibited longer steps on the side of injection, which caused contralateral turns (Figures 5E and 5F). For each animal, we observed a positive correlation between the difference in stride length and heading direction (*r* = 0.34-0.53). Together these data lead to a model where the difference in stride length on the left and right sides predicts locomotor direction (*r* = 0.96 for absolute distance, Figure 5E; *r* = 0.96 for log_2_ ratio, Figure 5F). These data are compatible with the hypothesis that increased *Chx10* Gi activity reduces locomotor rhythmogenic activity (9) on the ipsilateral side, leading to shorter steps on the side of the turn.

### *Chx10* Gi neurons function to control left/right movements during natural exploratory behaviors

To test the necessity/sufficiency of *Chx10* Gi neurons for turns during natural exploratory behaviors, we designed a paradigm where animals explored a novel left- or right-turned maze (Figure 6A). Animals were placed in the center of this simple spiral-shaped maze, which they explored until they exited the maze or until 10 minutes had elapsed. Unaffected mice rapidly completed both left- and right-turned mazes (Figures 6A-6D). In contrast, unilateral activation of *Chx10* stop neurons with hM3Dq-DREADDs increased exploratory time in a contralateral but not ipsilateral maze (Figures 6A and 6B), with several mice failing to complete the contralateral maze altogether (*n* = 6/8, Figure 6B). Mirroring this effect, inactivation of *Chx10* stop neurons with TeLC increased exploratory time in an ipsilateral but not contralateral maze (Figure 6C), with several mice failing to complete the ipsilateral maze (*n* = 3/8, Figure 6D). These experiments show that animals do not have an ability to compensate for dysfunction of the *Chx10* turning system, which appears to be required for natural exploratory function.

**Figure 6.**
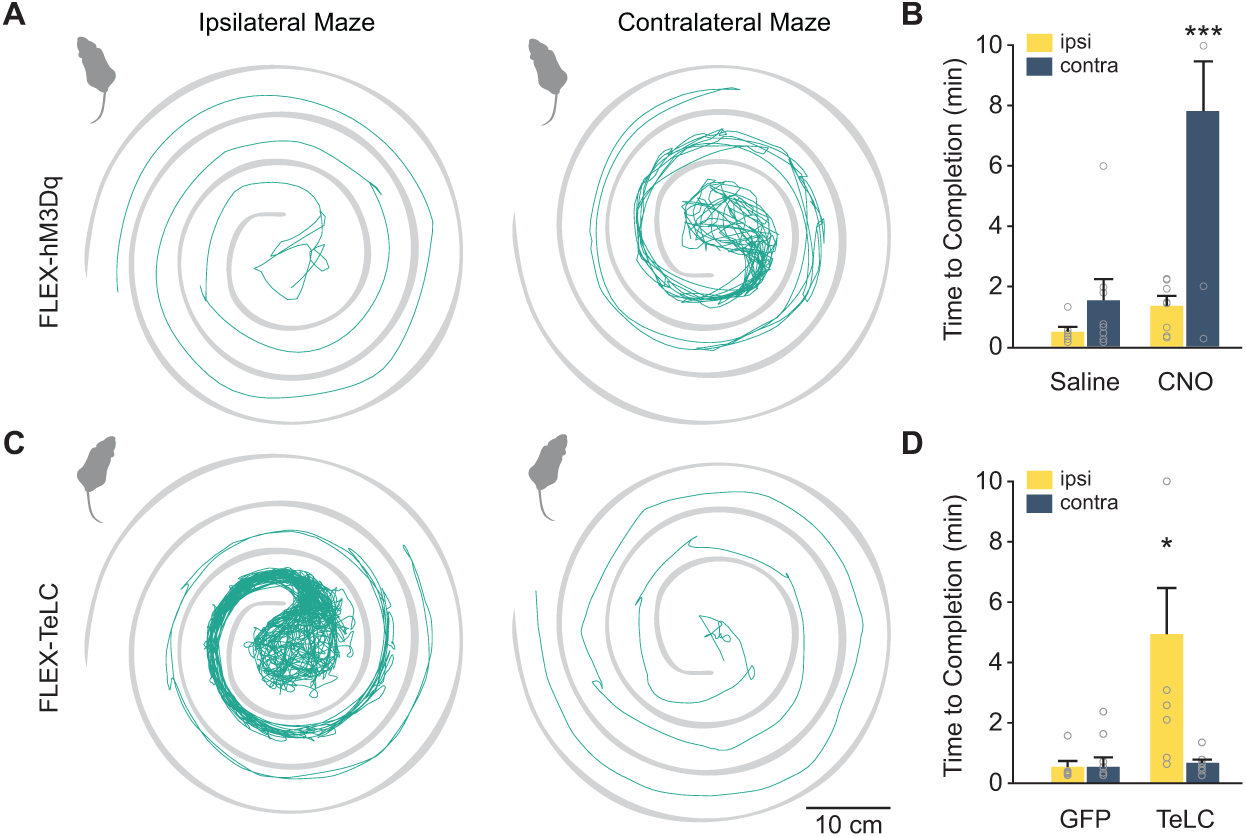
Unilateral function of *Chx10* Gi neurons is critical for left/right movements during natural exploratory behaviors. **A**, Activation of *Chx10* Gi neurons unilaterally with excitatory hM3Dq-DREADDs impairs exploration in a contralateral (right-turn) maze. **B**, 6 of 8 CNO injected animals failed to complete the contralateral maze during a 10 min trial. \*\*\**P* < 5×10^−5^, one-way ANOVA with Tukey HSD, *n* = 8. **C**, Inhibition of *Chx10* Gi neurons unilaterally with TeLC impairs exploration in an ipsilateral (left-turn) maze. **D**, 3 of 8 TeLC injected animals failed to complete the ipsilateral maze during a 10 min trial. \**P* < 0.011, one-way ANOVA with Tukey HSD, *n* = 7 for GFP and *n* = 8 for TeLC.

### Neurons of the superior colliculus provide a unilateral input that links *Chx10* Gi neurons to natural left/right movements

These experiments define a final command pathway for executing left/right locomotor asymmetries. The remaining question is whether it is possible to recruit *Chx10* Gi neurons unilaterally during natural behaviors? To address this question, we first performed monosynaptically restricted transsynaptic labeling from *Chx10* Gi neurons to determine their immediate synaptic inputs (47, 48). Cre-dependent rabies helper virus (AAV-FLEX-helper, see methods) was injected in the left Gi of *Chx10*^*Cre*^ mice, followed by RΔVG-4mCherry(EnvA). Consistent with our results using anterograde tracing (Figure 1B), monosynaptic retrograde tracing demonstrated that *Chx10* Gi neurons do not exhibit strong connectivity across the midline (Figures S4B and S4C); *Chx10* Gi neuronal populations are not reciprocally connected. *Chx10* Gi neurons received several long-range inputs that were primarily unilateral from the contralateral superior colliculus (SC), the ipsilateral zona incerta, the ipsilateral mesencephalic reticular formation (mRt) and the contralateral medial cerebellar (the fastigial) nucleus (Figures 7A and S4, Table S1). *Chx10* Gi neurons also exhibited bilateral long-range inputs from the lateral cerebellar (the dentate) nuclei and sensorimotor cortex (Figure S4, Table S1). Notably, no transsynaptic labeling was observed from the cuneiform or pedunculopontine nuclei, which comprise the midbrain locomotor region (Table S1). These experiments provide anatomical evidence that activity of *Chx10* Gi neurons can be biased via unilateral input from upstream nuclei.

**Figure 7.**
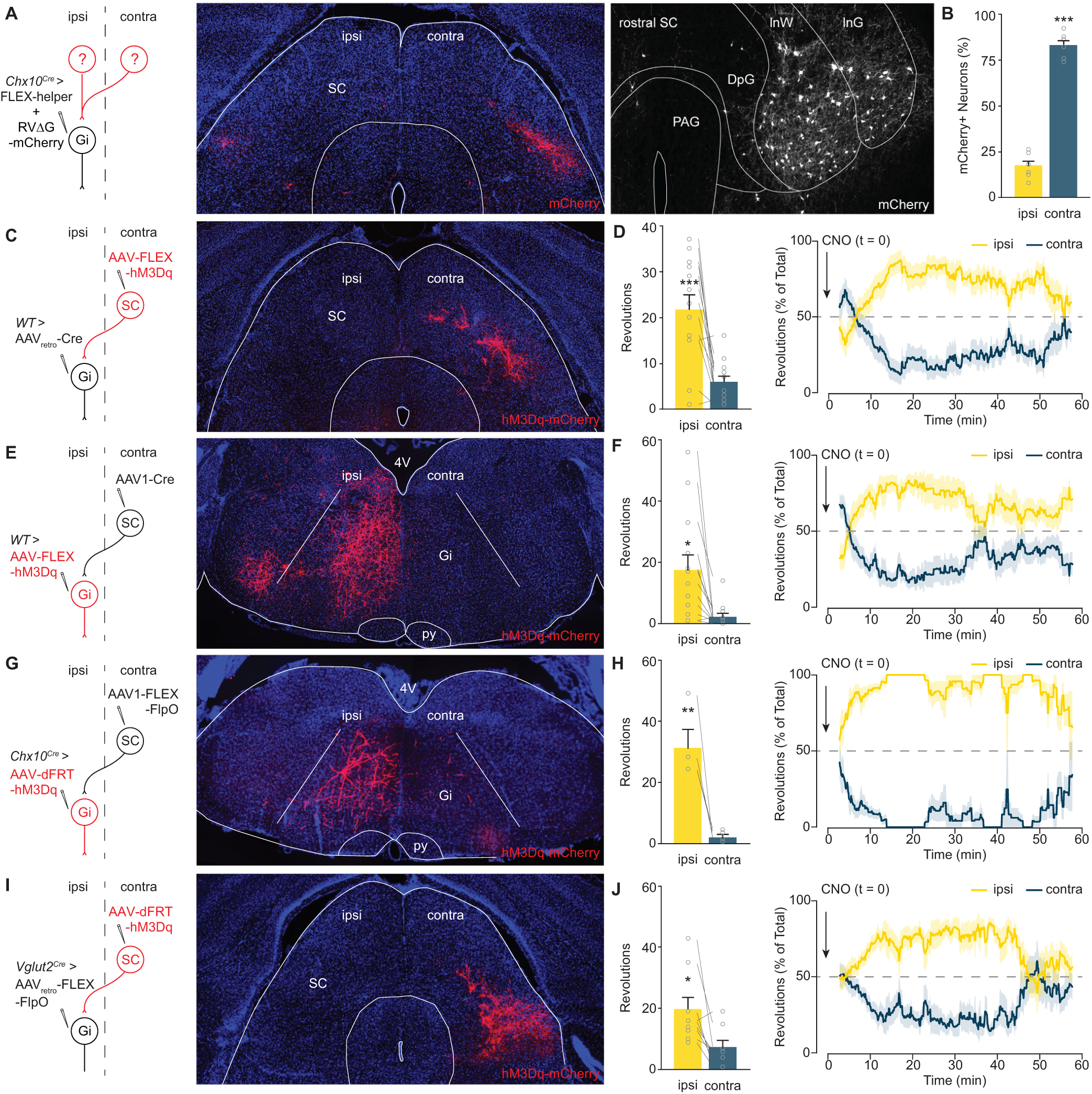
Contralateral SC steering acts through *Chx10* Gi neurons. **A**, A rabies transsynaptic tracing approach was used to identify presynaptic inputs to *Chx10* Gi neurons (see also, Figure S4 and Table S1). Rabies-mCherry tracing revealed a prominent input from the contralateral SC, where neurons occupied intermediate layers (*left*, caudal SC; *right*, rostral SC). **B**, Quantification of mCherry-labeled neurons in the ipsilateral and contralateral SC. \*\*\**P* = 1.8×10^−8^, two-tailed t-test, *n* = 6. **C**, Retrograde behavioral interrogation of the SC-Gi synapse in wild type mice. Injection of AAV_retro_-Cre in the Gi caused recombination of a FLEX-hM3Dq-mCherry virus injected in SC_contra_. **D**, CNO injection caused an ipsilateral turning preference, \*\*\**P* = 9.2×10^−5^, two-tailed t-test, *n* = *13. Right*, Instantaneous quantification of turning percentage after CNO injection. **E**, Anterograde behavioral interrogation of the SC-Gi synapse in wild type mice. Injection of AAV1-Cre in SC_contra_ caused recombination of a FLEX-hM3Dq-mCherry virus injected in Gi_ipsi_. **F**, CNO injection caused an ipsilateral turning preference. \**P* = 0.011, two-tailed t-test, *n* = 13. *Right*, Instantaneous quantification of turning percentage after CNO injection. **G**, Anterograde behavioral interrogation of the SC-Gi synapse in *Chx10*^*Cre*^ mice. Injection of AAV1-FLEX-FlpO in SC_contra_ caused recombination of a dFRT-hM3Dq-mCherry virus injected in Gi_ipsi_. **H**, CNO injection caused an ipsilateral turning preference. \*\**P* = 0.003, two-tailed t-test, *n* = 4. Right, Instantaneous quantification of turning percentage after CNO injection. **I**, Retrograde behavioral interrogation of the SC-Gi synapse in *Vglut2*^*Cre*^ mice. Injection of AAV_retro_-FLEX-FlpO in Gi_ipsi_ caused recombination of a dFRT-hM3Dq-mCherry virus injected in SC_contra_. **J**, CNO injection caused an ipsilateral turning preference. \**P* = 0.016, two-tailed t-test, *n* = 9. *Right*, Instantaneous quantification of turning percentage after CNO injection.

To test this possibility we stimulated the SC-Gi projection. We focused on the SC because of its prominent role in sensorimotor integration (49–51), and for its role in gaze and head orientation within 3D space (52). Projections from the contralateral SC to Gi have been described (51). We now show that these SC-Gi projection neurons occupy intermediate layers of the SC, and they synapse specifically with *Chx10* Gi neurons (Figures 7A and 7B). We tested the behavioral significance of this SC-Gi projection in four different ways. First, the SC-Gi projection was targeted by retrograde labeling of SC neurons using Gi_ipsi_ > AAV_retro_-Cre followed by SC_contra_ > FLEX-hM3Dq (Figure 7C). SC neurons targeted using this approach had a discrete laminar position which matched the rabies transsynaptic tracing (Figures 7A and 7C). In these mice, CNO administration rapidly initiated ipsilateral turning (Figure 7D). Next, postsynaptic Gi neurons were targeted using SC_contra_ > AAV1-Cre, an anterograde transsynaptic virus (51), followed by Gi_ipsi_ > FLEX-hM3Dq (Figure 7E). Gi neurons exhibited robust recombination on the ipsilateral side (Figure 7E). Again, CNO administration rapidly initiated ipsilateral turning (Figure 7F).

We supplemented these approaches using an intersectional strategy to allow both projection- and target-specific recombination in postsynaptic neurons (Figures 7G-J). In *Chx10*^*Cre*^ mice, targeting postsynaptic Gi neurons using SC_contra_ > AAV1-FLEX-FlpO followed by Gi_ipsi_ > dFRT-hM3Dq caused recombination in ipsilateral *Chx10* Gi neurons (Figure 7G). CNO administration rapidly initiated ipsilateral turning (Figures 7H). Finally, we showed that SC-Gi projection neurons are glutamatergic. In *Vglut2*^*Cre*^ mice (53), glutamatergic SC-Gi projection neurons were targeted using Gi_ipsi_ > AAV_retro_-FLEX-FlpO followed by SC_contra_ > dFRT-hM3Dq (Figure 7I). CNO administration rapidly initiated ipsilateral turning (Figure 7J). These results show that a natural sensorimotor pathway can bias *Chx10* Gi neuronal activity during spontaneous locomotion. Moreover, these experiments unambiguously demonstrate that the SC imparts left/right locomotor directional commands via *Chx10* Gi neurons.

## DISCUSSION

The present study uncovers a command system that enables left/right locomotor asymmetries necessary for directional movements. This asymmetric reticulospinal command system functions in parallel with symmetric start and speed control circuits (10, 11, 14, 18), and may be used symmetrically to arrest ongoing locomotion (9). Our study, therefore, completes the description of the three main requisite control components for locomotion: start, stop, and direction. Remarkably, when *Chx10* Gi neurons were biased to produce either ipsilateral or contralateral movements, animals could not perform a compensatory turn. This observation suggests that the turning system revealed here is the dominant system used during natural behaviors, and is recruited by brain circuits involved in meditating directional movements.

### Mechanism for locomotor turns

In aquatic vertebrates, left/right steering movements are achieved by asymmetric activity of glutamatergic reticulospinal neurons on the left and the right sides. This causes a forceful contraction of axial muscles on the side of the turn (54–59). In our experiments, we show that asymmetric movements in mice are also caused by an imbalance in descending excitation between the left and right sides. Nonetheless, the limb-based mechanism for turning is opposite to that described in species which use axial muscles as the predominant locomotor effector: Unilateral activation of mammalian *Chx10* Gi neurons initiates ipsilateral turning, which is caused by a reduction of locomotor speed and stride length on the ipsilateral side. Thus, for the limbs, excitation in spinal limb locomotor circuits is reduced on the side of the turn. In previous studies we showed that bilateral activation of *Chx10* Gi neurons arrests or reduces limbed locomotor activity via a brake on locomotor rhythm generation in the cord (9). The unilateral *Chx10* Gi gain- and loss-of-function experiments demonstrate that a unilateral brake (activation of *Chx10* Gi neurons) or release of the brake (inactivation of *Chx10* Gi neurons) is sufficient to mediate a limb-based turn—where speed and stride length are higher on the side opposite to the turn.

We suggest that this turning mechanism for limbed species evolved to account for general features of the limbed body plan. Chiefly, axial muscles generate force perpendicular to the directional axis. In contrast, limbs generate force parallel to the directional axis but at a distance from the midline, creating a moment about the medial axis. Although locomotor direction is ultimately mediated by the limbs and requires locomotion (60), unilateral stimulation of *Chx10* Gi neurons evoked bending of the head and trunk toward the side of stimulation even at rest. Based on these observations we propose that limbed animals use a two-component system for directing movements to the left or right side. Perhaps not surprisingly, this two-component biological turning system is a design principle adopted for steering four-wheeled vehicles millions of years after it was selected during evolution to control left/right locomotor asymmetries in quadrupeds: turning in quadrupeds and four-wheeled vehicles is enabled by a dedicated steering/differential system for independent control of speed on the left and right sides.

### Integrated function of *Chx10* Gi neurons

Mammalian *Chx10* Gi neurons were initially associated with locomotor stop, a behavioral response caused by bilateral activation (9). The data presented here give clear evidence that asymmetric engagement of *Chx10* Gi neurons causes turning. *Chx10* Gi neurons may therefore have a dual function—stop or turn—dependent on their symmetric or asymmetric activation. The basis for differential engagement may rest in task-dependent recruitment from upstream brain areas. Accordingly, our rabies tracing screen demonstrated that certain presynaptic nuclei contribute prominent bilateral or unilateral input. Prominent bilateral inputs from the lateral deep cerebellar nuclei and sensorimotor cortex (among others) provide an anatomical basis for a symmetric stop command. Prominent unilateral input from glutamatergic neurons in the intermediate area of the contralateral superior colliculus, the ipsilateral zona incerta, the ipsilateral mRt, and the contralateral medial deep cerebellar (fastigial) nucleus provide a basis for recruiting an asymmetric turn command. Indeed, we demonstrate that activation of the crossed SC-*Chx10* Gi pathway mediates movements contralateral to the superior colliculus. These experiments clearly demonstrate that *Chx10* Gi neurons can be recruited to generate locomotor asymmetries through the superior colliculus, a hub for sensorimotor integration (51, 52).

Interestingly, the superior colliculus, zona incerta, and mRt were recently identified in a brain-wide screen for neurons which control the decision to move left or right (31). Further, it has been shown that glutamatergic fastigial neurons project to the contralateral Gi (61). The present brain-wide screen links this excitatory cerebellar output to *Chx10* Gi neurons. Fastigial neurons are downstream of the vestibule-cerebellum, which has been shown to encode turning during locomotion (62). Glutamatergic inputs from contralateral SC and fastigial neurons to *Chx10* Gi neurons, therefore, contribute information from both sensorimotor and vestibular sources, information which ultimately biases locomotor movements to the ipsilateral side.

Together our findings suggest that *Chx10* Gi neurons act as the final common effector for left/right movements, and might act as a locus for integration of diverse sensory signals which contribute to the decision to move left or right. Understanding the specific contribution of brain areas upstream of *Chx10* Gi neurons—and how they work in concert—is expected to lead to a more thorough understanding of how motor control is organized at the behavioral level.

## Acknowledgments

We thank K. Sharma, L. Zagoraiou, S. Crone, and T.M. Jessell for the *Chx10*^*Cre*^ mouse. We acknowledge the Core Facility for Integrated Microscopy, Faculty of Health and Medical Sciences, University of Copenhagen. We thank Iryna Vesth-Hansen, Dorthe Meinertz, and Paulina Wanken for technical assistance, and members of Ole Kiehn’s lab for discussion and comments on previous versions of this manuscript. This work was supported by an EMBO Long-Term Fellowship (J.M.C., ALTF 421-2018), a European Research Council Advanced Grant (O.K., LocomotorIntegration, REP-SCI-693038), and the Novo Nordisk Laureate Program (O.K., NNF15OC0014186).

## Author contributions

Conceptualization, J.M.C. and O.K.; Methodology, J.M.C., R.L., A.M., and O.K.; Investigation, J.M.C., R.L., A.M.; Resources – I.R.W.; Writing – Original Draft, J.M.C., O.K.; Writing – Review & Editing, J.M.C., R.L., and O.K.; Supervision – O.K.; Funding Acquisition, J.M.C., O.K.

## MATERIALS AND METHODS

### Mice

All animal experiments and procedures were approved by Dyreforsøgstilsynet in Denmark and the local ethics committee at the University of Copenhagen. The *Chx10*^*Cre*^ mouse is the same as that used previously (9, 63). The *Vglut2*^*Cre*^ mouse is described in (53). *R26R*^*ChR2-EYFP*^ mice were obtained from Jackson Laboratories (Jackson Stock 012569). For *in vitro* experiments, we used newborn mice from *Chx10*^*Cre*^ and *R26R*^*ChR2-EYFP*^ crosses. For *in vivo* experiments, we used hemizygous *Chx10*^*Cre*^, *Vglut2*^*Cre*^, or wild type mice (C57BL6/J (Jackson Stock 000664) greater than 8 weeks of age. Experiments were performed with similar numbers of male and female mice.

### *In Vitro* Recording and Optogenetics

Locomotor-like activity was recorded from *in vitro* brainstem-spinal cord preparations isolated from P0-4 *Chx10*^*Cre*^;*R26R*^*ChR2-EYFP*^ mice. Neonatal mice were anesthetized, and decapitated at the level of the midbrain. Mice were then eviscerated, and vertebral bodies were quickly removed ventrally from the level of the rostral pons caudally to the sacral spinal cord under ice-cold oxygenated dissection buffer (95% O_2_/5% CO_2_, 4 °C) composed of 111 mM NaCl, 3 mM KCl, 26 mM NaHCO_3_, 1.1 mM KH_2_PO_4_, 0.25 mM CaCl_2_, 3.7 mM MgCl_2_, and 11 mM D-glucose. The ventral roots of the lumbar spinal cord were cut at their point of exit from the vertebral canal. The caudal neuroaxis was then isolated from the vertebral canal, and a coronal section was performed at the level of the facial motor nucleus (VII), rostral to the anterior inferior cerebellar artery. The rostral aspect of the preparation was maintained in an upward position to allow access for optical stimulation (9). Alternatively, the brainstem was split along the midline from the most rostral part to C1. The optical stimulation was then performed unilaterally from the cut surface.

A two-compartment system was used for pharmacological separation of the brainstem and lumbar spinal cord. The preparation was pinned to a Sylgard stage, and Vaseline was applied to the upper thoracic spinal cord at T8. A peristaltic pump was used to separately perfuse the rostral (brainstem) and caudal (lumbar spinal cord) compartments with oxygenated recording buffer (95% O_2_/5% CO_2_, 22-24 °C), composed of 111 mM NaCl, 3 mM KCl, 26 mM NaHCO_3_, 1.1 mM KH_2_PO_4_, 2.5 mM CaCl_2_, 1.25 mM MgCl_2_, and 11 mM D-glucose. Kynurenic acid (KA, 4 mM) was applied to the brainstem compartment to block all glutamatergic transmission in the brainstem thereby isolating the contribution of *Chx10* reticulospinal neurons to spinal circuits (9). A cocktail of serotonin (5HT, 8 µM) and N-methyl-D-aspartic acid (NMDA, 8 µM) was applied to the lumbar spinal cord compartment to initiate locomotor-like activity. Fast green dye was maintained within the rostral compartment to verify an intact diffusion barrier, that is, drugs from the rostral compartment did not mix with the caudal compartment and *vice versa*.

Bipolar suction electrodes were attached to the right and left L2 ventral roots. The signal was amplified 5000-10000 times, band-pass filtered from 100 Hz to 1 kHz and sampled at a frequency of 1 kHz.

Optogenetic stimulation of ChR2-expressing neurons was performed using a 473 nm laser system (UGA-40; Rapp Opto-electronic), which delivered blue light at an intensity of 30 mW mm^−2^ (64). Blue light was directed at the preparation using an optical fiber (200 µm core, 0.22 NA, Thorlabs). Photo illumination was carried out continuously for 5-15 s. Burst amplitude (Figure 2D) was analyzed from rectified, integrated (bin = 0.25 s) traces, where we quantified 5 bursts before, those bursts during, and 5 bursts after light stimulation. 1-6 trials were quantified for each animal.

### Stereotaxic Injections

Viral injections in adult mice were performed using a stereotactic injection system (Neurostar, Tübingen, Germany). Mice were anesthetized with 4% isofluorane, and maintained under anesthesia with 2% isofluorane for the duration of the surgery. Anesthetic depth was verified using a toe-pinch test. Virus was mixed with fast green for visualization, and injected using a glass micropipette at a rate of 100 nl min^−1^. The glass micropipette was held in place for 5 min following injection to prevent backflow.

For unilateral anterograde tracing from Gi to the spinal cord, we injected 250 nl AAV1-phSyn1(S)-FLEX-tdTomato-T2A-SypEGFP-WPRE (5.56×10^11^ ml^−1^, Viral Vector Core, Salk Institute for Biological Sciences; Addgene 51509) (65) in *Chx10*^*Cre*^ animals at least 8 weeks of age. Mice were perfused 6 weeks after the injection. All injections in Gi were made at 6.0 mm AP, ± 0.8 mm ML, and 5.5 mm DV relative to bregma.

For experiments using excitatory hM3Dq-DREADDs, *Chx10*^*Cre*^ animals at least 8 weeks of age were injected in either the left or right Gi with 500 nl of AAV5-hSyn1-DIO-hM3D(Gq)-mCherry-WPRE (6×10^12^ ml^−1^, Viral Vector Facility, University of Zurich, v89) or 500 nl AAV5-hSyn1-DIO-mCherry control virus (1.3×10^13^ ml^−1^, Viral Vector Facility, University of Zurich, v84). Experiments were performed 3-6 weeks after injection. For *in vivo* optogenetics experiments, *Chx10*^*Cre*^ animals were injected with 500 nl of AAVdj-Ef1a-DIO-hChR2(E123T/T159C)-P2A-mCherry-WPRE in the left or right Gi. Optical fibers were implanted 3 weeks later at −6 mm AP, ± 0.5 mm ML, −4.8 mm DV relative to bregma, and photostimulation experiments were performed the next day. For experiments using tetanus toxin virus, we injected either 300 or 500 nl of AAV1-FLEX-TeLC-EGFP to inhibit *Chx10* Gi neurons (9, 66), or 500 nl AAV5-FLEX-EGFP as a control. Tetanus toxin behavioral experiments were performed within 9 days of viral injection. For inhibitory DREADDs, *Chx10*^*Cre*^ mice were injected with 350 nl of AAV-FLEX-hM4Di-mCherry in the left or right Gi (7.4×10^12^ ml^−1^, Viral Vector Facility, University of Zurich, v84). Experiments were performed 3-6 weeks after injection.

For rabies transsynaptic labeling, the left Gi was injected with 200 nl of a 1:1 mixture of AAV-syn-FLEX-splitTVA-EGFP-tTA (Addgene 100798) and AAV-TREtight-mTagBFP2-B19G (Addgene 100799) (**?**). Seven days later, we injected 500 nl of RΔVG-4mCherry(EnvA) in the same location. Mice were perfused 7 days after the second injection. The synthesis of pAAV-syn-FLEX-splitTVA-EGFP-tTA, pAAV-TREtight-mTagBFP2-B19G, and RΔVG-4mCherry(EnvA) has been described (47, 67).

For targeting the SC_contra_-Gi_ipsi_ projection, injection of AAV-DIO-hM3Dq-mCherry or AAV5-dFRT-hM3Dq-mCherry (3.5×10^12^ ml^−1^, Viral Vector Facility, University of Zurich, v189-5) was followed 1 week by injection of AAV_retro_-EGFP-Cre (1.3×10^13^ ml^−1^, Addgene, 105540-AAVrg), AAV_retro_-FLEX-EGFP-2A-FlpO (6.8×10^12^ ml^−1^, Viral Vector Facility, University of Zurich, v171-retro), AAV1-EGFP-Cre (1×10^13^ ml^−1^, Addgene, 105540-AAV1), or AAV1-FLEX-EGFP-2A-FlpO (7.3×10^12^ ml^−1^, Viral Vector Facility, University of Zurich, v171-1). Behavioral experiments were performed 1-2 weeks following second viral injection. For retrograde targeting of SC_contra_-Gi_ipsi_ projection neurons in wild-type mice, 80 nl of AAV-DIO-hM3Dq-mCherry was injected in SC_contra_ and 150 nl of AAV_retro_-EGFP-Cre in Gi_ipsi_. For anterograde targeting of SC_contra_-Gi_ipsi_ postsynaptic neurons in wild-type mice, 40 nl of AAV1-EGFP-Cre was injected in SC_contra_ and 150 nl of AAV-DIO-hM3Dq-mCherry in Gi_ipsi_. For retrograde targeting of SC_contra_-Gi_ipsi_ projection neurons in *Vglut2*^*Cre*^ mice, 80 nl of AAV-dFRT-hM3Dq-mCherry was injected in SC_contra_ and 150 nl of AAV_retro_-FLEX-EGFP-2A-FlpO in Gi_ipsi_.For anterograde targeting of SC_contra_-Gi_ipsi_ *Chx10* postsynaptic neurons in *Chx10*^*Cre*^ mice, 80 nl of AAV1-FLEX-EGFP-2A-FlpO was injected in SC_contra_ and 500 nl of AAV-dFRT-hM3Dq-mCherry in Gi_ipsi_. SC coordinates used for these experiments were based on localization of RVΔG-4mCherry(EnvA) labeled cells (68): SC: −3.5 mm AP, ± 1.12 mm ML, and 2.15 mm DV relative to bregma.

### *In Vivo* Optogenetics

Optical fibers (200 µm core, NA 0.22, Thorlabs) were implanted 3 weeks after ChR2 viral injection (see information on injection/implantation above). Photostimulation experiments were carried out the day after implantation. Fibers were held in a 1.25 mm ferrule, and coupled to a 473 nm laser (Optoduet, Ikecool Corporation) via a ceramic mating sleeve. Photostimuli were manually triggered via a TTL-pulse given by Ethovision to a Master-8 pulse generator. Laser power was adjusted at 40 Hz (10 ms pulses for 1 s total duration) to initiate a strong turning response in each animal. We found that the laser power necessary to initiate a response was graded between 5-20 mW.

### Limb Dynamics

DeepLabCut (45, 69) was used for markerless extraction of paw position in an openfield. DeepLabCut 2.0 was installed on a PC equipped with a GEforce RTX 2080 Ti graphics card. We captured videos of mice moving from below (50 f.p.s.). Body center position was tracked using Ethovision XT, and video segments were extracted for locomotor bouts directed through the center of the arena that were >5 cm s^−1^ and >15 cm long. The four paws, the tip of the nose, and the base of the tail were tracked using DeepLabCut 2.0. 200 frames were randomly selected for labeling. The network was trained for 200,000 iterations until the loss reached a plateau. Segments of recording with likelihood <0.6 for all markers were excluded.

Subsequent analyses of limb kinematics were made using custom scripts in Python 3.7. The body angle was defined as tail base-nose angle relative to the x axis. The velocity of each paw was used to define steps. The beginning of the swing phase was defined as the point where paw velocity passed above 9 cm s^−1^, and the end of the swing phase (i.e., the beginning of the stance phase) was defined as the point where paw velocity passed back below 9 cm s^−1^. During alternating gates including fast walk and trot, animals move forward through the environment using diagonals—i.e. one forelimb and the contralateral hindlimb. Diagonal steps were defined as those occurring near simultaneously (within 0.1 s) for diagonal limbs. For each paw, stride length was calculated as the length of the segment between the paw at the beginning and end of the diagonal step. We compared differences in stride length between the left and right sides with body angle at the beginning and end of a diagonal step. Pearson’s correlation coefficient was used to estimate how stride length correlated with body angle.

### Tissue Immunochemistry & Imaging

Mice were euthanized by anesthetic overdose with pentobarbital (250 mg kg^−1^), and perfused transcardially with 4 °C saline followed by 4% paraformaldehyde. Brain and spinal cord tissue was dissected free, and then post-fixed in 4% paraformaldehyde for 3 h at 4 °C. Tissue was cryoprotected by incubation in 30% sucrose in phosphate buffered saline (PBS) overnight. Tissue was then embedded in Neg-50 medium (ThermoFisher Scientific) for cryostat sectioning. Coronal or sagittal sections were obtained on a Leica cryostat and mounted on Superfrost Plus slides (ThermoFisher Scientific). Brainstem coronal sections were cut at 30 µm thickness, whereas spinal cord coronal sections were cut at 20 µm. Sagittal sections were cut at 30 µm.

Sections were rehydrated for 5 minutes in PBS + 0.5% Triton-X100 (PBS-T; Sigma-Aldrich), and then blocked for 2 h in 10% normal donkey serum in PBS-T (Jackson ImmunoResearch). Sections were incubated overnight with primary antibodies diluted in blocking solution. We used the following primary antibodies: chicken anti-GFP (1:1000, Abcam, ab13970) and rabbit anti-DsRed/tdTomato/mCherry (1:1000, Clontech, 632496). Slides were washed 4 times in PBS-T, and then incubated with appropriate donkey secondary antibodies diluted in blocking solution (1:500, ThermoFisher Scientific). Slides were washed 4 times in PBS-T, counterstained with Hoechst 33342 (1:2000) or NeuroTrace 435 (1:400, ThermoFisher Scientific), and were mounted with coverslips using mowiol 4-88 medium. Sections were imaged using either a Zeiss widefield epifluorescence microscope or a Zeiss LSM 780 confocal microscope.

### Drugs

Clozapine-N-oxide (CNO, Tocris, 4936) was dissolved in saline immediately prior to behavioral experiments. CNO was administered intraperitoneally at a dose of 0.5 mg kg^−1^ (*Chx10*^*Cre*^ > FLEX-hM3Dq; Figure 3, Figure S2) or 1 mg kg^−1^ (*Chx10*^*Cre*^ > FLEX-hM4Di, Figure 4; SC or Gi > FLEX-hM3Dq or dFRT-hM3Dq, Figure 7).

### Cylinder Test

Mice were placed in a 15 cm diameter cylinder for 10 minutes. Ethovision XT (Noldus) software was used to capture video (15 f.p.s.) and carry out tracking of head, center point, and tail. 360° clockwise and counterclockwise revolutions were quantified (tail point to center point, 50° threshold).

### Open Field

Mice were acclimated to the behavioral suite prior to testing in an open field arena. Open field analysis was carried out in a 50 × 50 cm square arena illuminated with an infrared lamp. Ethovision XT (Noldus) software was used to capture video (15-25 f.p.s.) and carry out tracking analysis. Mice were tracked using head, center, and tail points, enabling quantification of rotations as a function of time. Movement parameters taken in the open field included total movement (m), ambulation (time moving at a velocity greater than 2 cm s^−1^), and 360° revolutions (tail point to center point, 50° threshold). Tracking was carried out for 60 minutes. Mice were allowed to acclimate to the arena for 30 min, followed by a 30 min probe. Movement parameters are reported for minutes 30-60.

### Ipsilateral/Contralateral Spiral Maze

Spiral-shaped mazes were used to test clockwise/counterclockwise movement preference. For this purpose, mazes were fabricated which could be explored from the center-point to the periphery only by moving in a clockwise or counterclockwise direction (see Figures 6A and 6C for illustration). These mazes exhibit an increasing radius as mice move from the center-point to the periphery. For each mouse, a 10 minute trial was conducted in either the left- or right-turned maze. One affected mouse exhibited a turning radius that was smaller than the center of the maze (see Figures 6A and 6C). This affected mouse was excluded from analysis because it could not complete either the ipsilateral or contralateral maze. Body tracking was carried out with Ethovision XT (Noldus) using center-point detection.

### Analysis

For quantification of viral transduction in Gi, the number of neurons ipsilateral and contralateral to the injection site was counted in every 10th section. These counts were used to estimate the percentage of viral transduction contained to the ipsilateral side (Figure 1). Similar analysis was performed to estimate the percentage of labeled neurons in the ipsilateral versus contralateral superior colliculus (Figure 7).

For analysis of synGFP^+^ punctae in tdTom-2A-synGFP traced projections, high resolution images of GFP-stained coronal spinal cord sections were obtained with a confocal microscope. Three images from each spinal level (i.e. cervical, thoracic, lumbar) were quantified from three mice. Analysis of synGFP punctae was restricted to the gray matter, with the assumption that synGFP^+^ punctae in white matter reflected transport of synGFP protein rather than *bona fide* synapses. SynGFP^+^ punctae were isolated from the images using thresholding, and the position of each punctum was extracted using the particle analysis feature in ImageJ (https://imagej.nih.gov/ij/). The position of the central canal was defined using the Nissl counterstain, and synGFP^+^ punctum position was normalized between sections and animals using this point as a reference. From this data set, the percentage of synapses ipsilateral and contralateral to the injection site was estimated (Figure 1). Synaptic density plots were constructed in R, where gradation represents the range from zero to maximum density at each spinal level (Figure 1G). Synaptic density plots represent the average of 3 animals. Coordinates and abbreviations are based on Paxinos and Franklin’s reference atlas (68), and corresponding plates are from the Allen Mouse Brain Atlas (70).

### Statistics

A two-tailed t-test was performed for pairwise comparisons. For multiple comparisons, a one-way ANOVA was performed to determine whether significant differences existed between conditions, followed by Tukey-Kramer HSD for assigning levels of significance. All data are given as mean ± standard error mean. Individual data points are plotted for every analysis. For each comparison, information on statistical analyses, *P* value statistics, and *n* values are listed in the figure legend. *n* values represent distinct biological replicates. *P* < 0.05 was considered statistically significant, where \**P* < 0.05, \*\**P* < 0.01, and \*\*\**P* < 0.001.

## Supplementary Material

**Figure S1.**
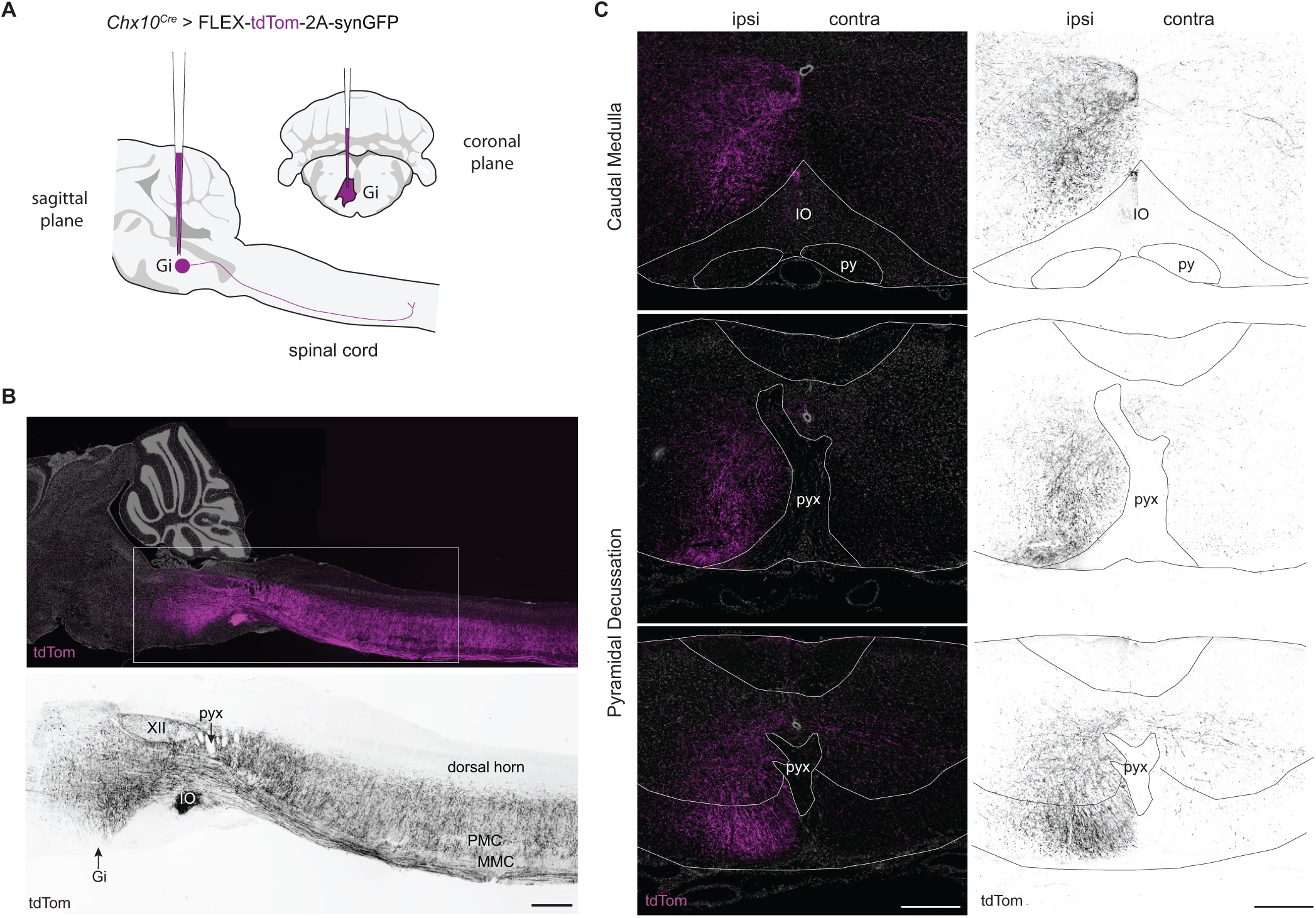
*Chx10* Gi neurons form a prominent tract of ipsilaterally projecting axons. **A**, Unilateral labeling of *Chx10* neurons of the rostral gigantocellularis (Gi) using the Cre-dependent anterograde tracer AAV-FLEX-tdTom-2A-synGFP. **B**, *Top*, Sagittal section of tdTom^+^ projections ipsilateral to the injection site. tdTom^+^ axons formed a prominent tract that projected caudally to the spinal cord. *Bottom*, inset from Top. Scale bar = 500 µm. XII, hypoglossal motor nucleus; pyx, pyramidal decussation; IO, inferior olive; PMC, phrenic motor column; MMC, medial motor column. **C**, Coalescence of *Chx10* reticulospinal axons dorsal to the inferior olive (top), and subsequent positioning in the ventrolateral funiculus at the level of the pyramidal decussation (bottom).

**Figure S2.**
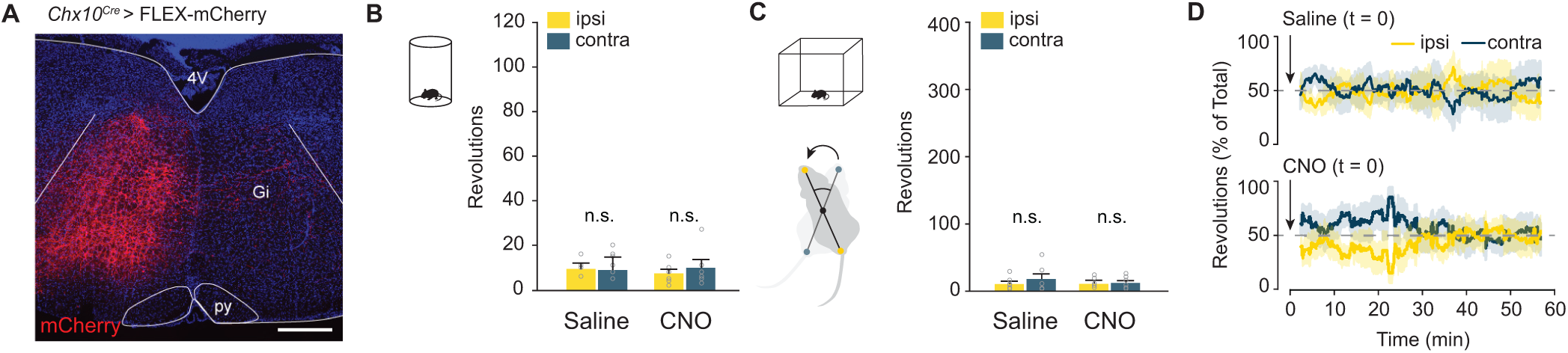
CNO administration does not affect turning preference in control AAV-mCherry injected mice. **A**, Example of mCherry expression 3 weeks after a 500 nl unilateral injection of AAV-FLEX-mCherry in a *Chx10*^*Cre*^ mouse. mCherry expression is confined to the side ipsilateral to injection. **B**, Turning preference in a 10 minute cylinder assay is unaffected by administration of CNO. *P* > 0.45, one-way ANOVA with Tukey HSD, *n* = 6. **C**, Turning preference in an open field arena is unaffected by administration of CNO. *P* > 0.77, one-way ANOVA with Tukey HSD, *n* = 6. **D**, Instantaneous quantification of ipsilateral and contralateral revolutions after injection of either saline or CNO in AAV-FLEX-mCherry injected mice.

**Figure S3.**
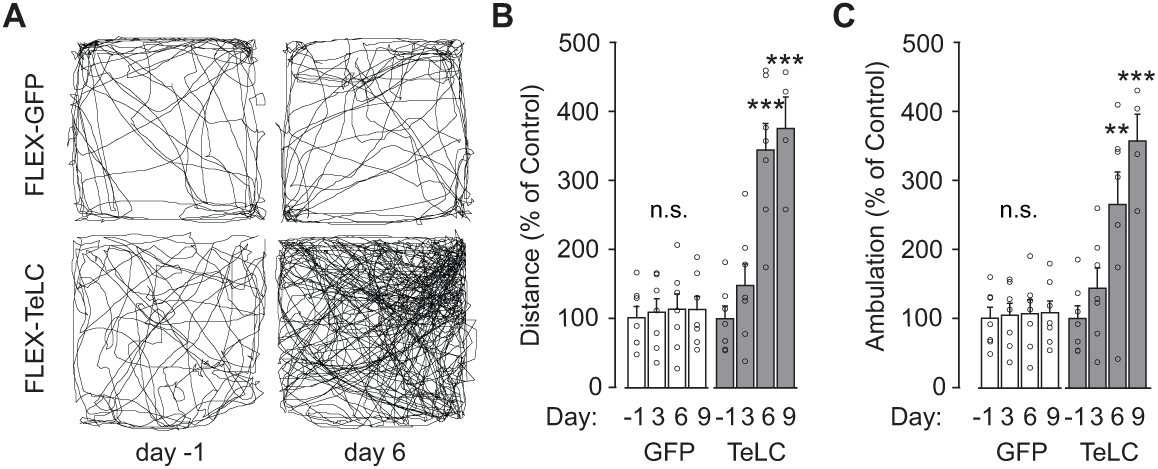
Unilateral silencing of *Chx10* Gi neurons increases locomotion in an open field assay. **A**, Open field analysis of a *Chx10*^*Cre*^ > FLEX-GFP or FLEX-TeLC mouse one day before AAV injection (day -1) and 6 days after AAV injection. Traces represent 15 minutes of open field activity. **B**, Total distance traveled during a 30 minute open field trial -1, 3, 6, or 9 days after AAV injection as a percentage of day -1. GFP: *P* > 0.97 for each comparison, *n* = 7. TeLC: \*\**P* < 1.2×10^−4^ (day 6 and 9 vs. -1), *n* = 7 for days -1, 3, and 6, *n* = 4 for day 9. One-way ANOVA with Tukey HSD. **C**, Open field ambulation as a percentage of day -1. GFP: *P* > 0.98 for each comparison. TeLC: \*\**P* = 0.009, day 6 vs. -1; \*\*\**P* = 6.6×10^−4^, day 9 vs. -1. One-way ANOVA with Tukey HSD.

**Figure S4.**
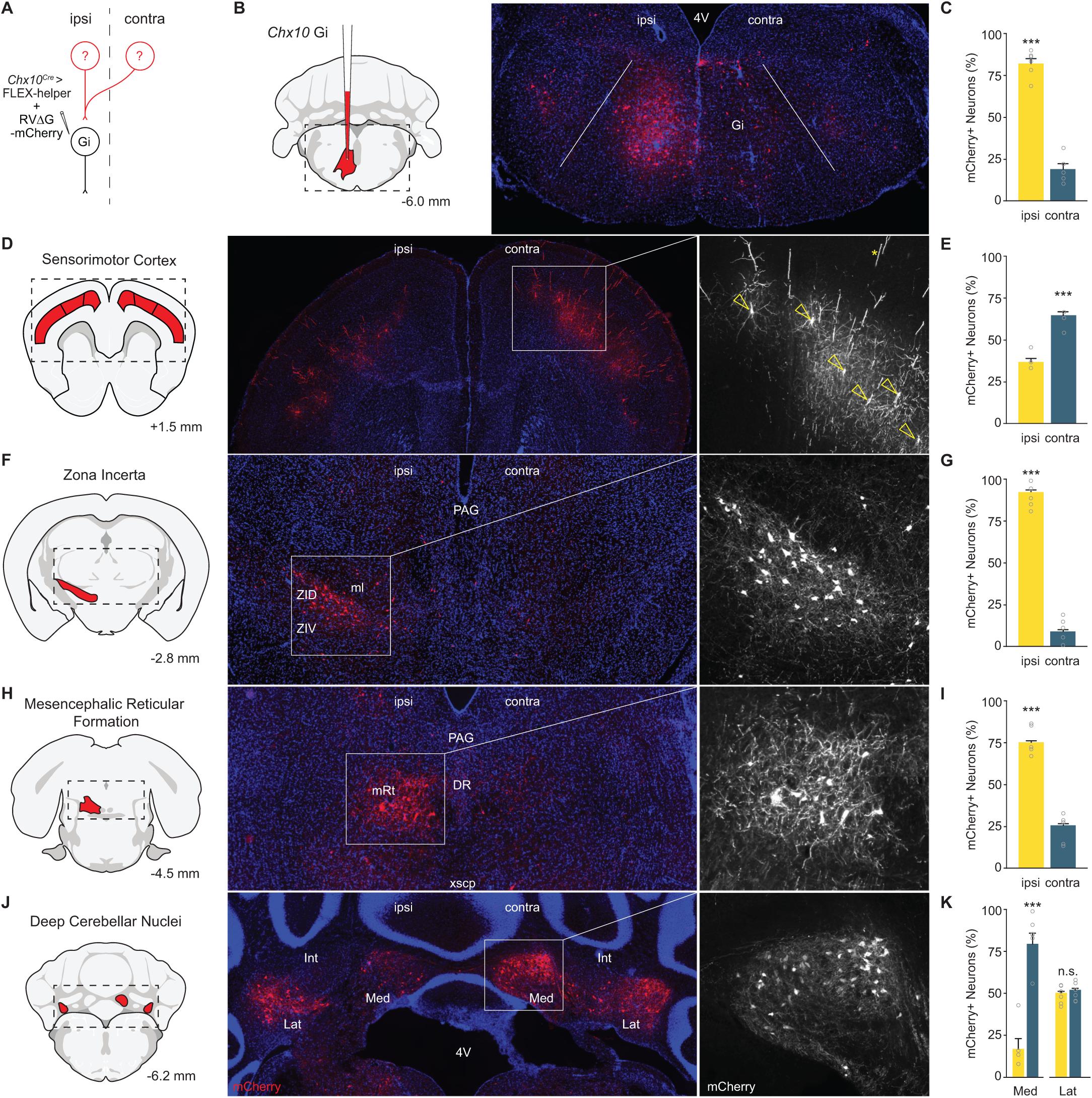
Monosynaptic rabies tracing identifies *Chx10* Gi presynaptic inputs. **A**, A rabies transsynaptic tracing approach was used to identify presynaptic inputs to *Chx10* Gi neurons (see also, Figure 7 and Table S1). **B**, Injection site in rostral Gi. Initial site of infection is visualized as a large population of mCherry^+^ neurons, accompanied by dense mCherry^+^ processes. Starter neurons of the ipsilateral Gi do not exhibit a strong input from neurons of the contralateral Gi. **C**, Quantification of mCherry-labeled neurons in the ipsilateral and contralateral Gi. \*\*\**P* = 2.5×10^−7^, two-tailed t-test, *n* = 6. **D**, Bilateral input to *Chx10* Gi neurons from neurons of primary motor and somatosensory cortex. Open yellow triangles point to soma from pyramidal neurons. Yellow asterisk indicates an apical dendrite. **E**, Quantification of mCherry-labeled neurons in the ipsilateral and contralateral cortex. \*\*\**P* = 1.9×10^−6^, two-tailed t-test, *n* = 6. **F**, Input to *Chx10* Gi neurons from neurons of the ipsilateral zona incerta. Presynaptic neurons were observed primarily in the dorsal (ZID) and caudal aspects of the zona incerta. **G**, Quantification of mCherry-labeled neurons in the ipsilateral and contralateral zona incerta. \*\*\**P* = 3.7×10^−9^, two-tailed t-test, *n* = 6. **H**, Unilateral input to *Chx10* Gi neurons from the ipsilateral mesencephalic reticular formation. **I**, Quantification of mCherry-labeled neurons in the ipsilateral and contralateral mesencephalic reticular formation. \*\*\**P* = 1.8×10^−6^, two-tailed t-test, *n* = 6. **J**, Medial (Med) and lateral (Lat) deep cerebellar nuclei exhibited unilateral or bilateral, respectively, input to *Chx10* Gi neurons. **K**, Quantification of mCherry-labeled neurons in the deep cerebellar nuclei. Med, \*\*\**P* = 1.1×10^−5^, two-tailed t-test, *n* = 6; Lat, *P* = 0.56, two-tailed t-test, *n* = 6.

**Table S1.**
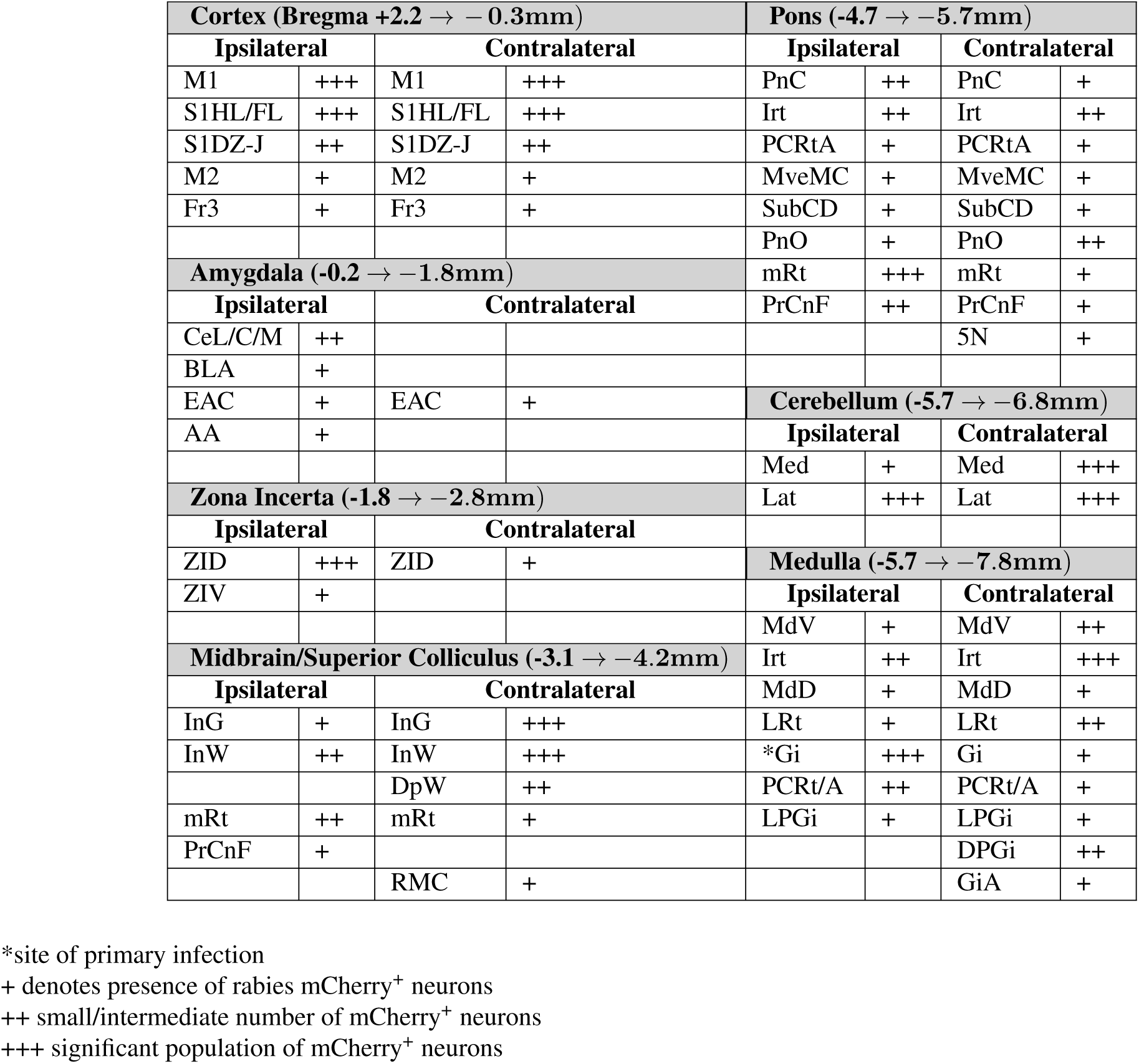
Summary of monosynaptic rabies tracing data.

See also, Figure 7 and Figure S4. Qualitative analysis is based on curation of rabies retrograde tracing data from *n* = 6 animals. Coordinates represent the rostrocaudal extent to which mCherry^+^ neurons were observed in each corresponding structure (e.g. Cortex, Amygdala, etc.). Coordinates and abbreviations are based on Paxinos and Franklin’s reference atlas (68).

Abbreviations: 5N, motor trigeminal nucleus; AA, anterior amygdaloid area; BLA, basolateral amygdaloid nucleus, anterior part; CeC/L/M, central amygdaloid nucleus, capsular/lateral//medial division; DPGi, dorsal paragigantocellular nucleus; DpW, deep white layer of the superior colliculus; EAC, extended amygdala, central part; Fr3, frontal cortex, area 3; Gi/A, gigantocellular reticular nucleus, alpha part; InG/W, intermediate gray/white layer of the superior colliculus; IRt, intermediate reticular nucleus; Lat, lateral cerebellar nucleus; LPGi, lateral paragigantocellular nucleus; LRt, lateral reticular nucleus; M1, primary motor cortex; M2, secondary motor cortex; MdD/V, medullary reticular nucleus, dorsal/ventral part; Med, medial cerebellar nucleus; mRt, mesencephalic reticular formation; MVeMC, medial vestibular nucleus, magnocellular part; PCRt/A, parvicellular reticular nucleus, alpha part; PnC/O, pontine reticular nucleus, caudal/oral part; PrCnF, precuneform area; RMC, red nucleus, magnocellular part; S1 DZ/J/HL/FL, primary somatosensory cortex, dysgranular zone/jaw region/hindlimb region/forelimb region, SubCD, subcoeruleus nucleus, dorsal part; ZID/V, zona incerta, dorsal/ventral part.

**Video S1. Open field analysis of a left Gi *Chx10***^***Cre***^ **> FLEX-hM3Dq targeted mouse 0, 15, and 25 min after administration of CNO**. Immediately following CNO administration (0 min), the mouse exhibits no preference in locomotor direction. After 15 min, the mouse exhibits a clear preference toward the ipsilateral side, making large left-turn circles in the arena. After 25 min, the mouse exhibits sharp rotations toward the ipsilateral side. Playback is three times real speed.

**Video S2. Photostimulation of left Gi *Chx10***^***Cre***^ **> FLEX-ChR2 targeted mice during spontaneous locomotion and at rest**.

**Video S3. Open field analysis of a left Gi *Chx10***^***Cre***^ **> FLEX-TeLC targeted mouse**. Before viral injection (day -1), the mouse exhibited no left or right preference in locomotor direction. Six days following viral injection, the mouse exhibits a clear locomotor preference toward the contralateral (right) side. Playback is three times real speed.

**Video S4. DeepLabCut tracking of footfalls and movement trajectories in wild-type, *Chx10***^***Cre***^ **> FLEX-hM3Dq, and *Chx10***^***Cre***^ **> FLEX-TeLC targeted mice**. Footfalls are plotted when the limb velocity reaches 0 cm s^−1^.

